# A novel self-organizing embryonic stem cell system reveals signaling logic underlying the patterning of human ectoderm

**DOI:** 10.1101/518803

**Authors:** George Britton, Idse Heemskerk, Rachel Hodge, Amina A Qutub, Aryeh Warmflash

## Abstract

During development, the ectoderm is patterned by a combination of BMP and WNT signaling. Research in model organisms has provided substantial insight, however, there are currently no systems to study this patterning in humans. Further, the complexity of neural plate border specification has made it difficult to transition from discovering the genes involved to deeper mechanistic understanding. Here, we develop an *in vitro* model of human ectodermal patterning, in which hESCs self-organize to form robust and quantitatively reproducible patterns corresponding to the dorsal-ventral axis of the embryo. Using this platform, we show that the duration of endogenous WNT signaling is a crucial control parameter, and that cells sense relative levels of BMP and WNT signaling in making fate decisions. These insights allowed us to develop an improved protocol for placodal differentiation. Thus, our platform is a powerful tool for studying human ectoderm patterning and for improving directed differentiation protocols.

## Introduction

The ectodermal germ layer arises as a spatially organized epithelium composed of progenitors of the neural plate, neural crest, placodes and epidermis along the dorsal-ventral axis of the developing embryo. The establishment of this pattern is required for neurulation in which the neural plate folds to form a tube with neural crest and placodal cells outside the tube at the dorsal end and covered by epidermis. Studies from model organisms have identified key morphogens and their inhibitors that coordinate the patterning of ectodermal fates. In several models, BMP4 was shown to induce placodal and epidermal fates ventrally (Kwon et al., 2010; Li et al., 2013; Reichert et al., 2013; Wilson et al., 1997), while neural crest induction requires WNT ligands secreted from the paraxial mesoderm or the ectoderm itself (Dorsky et al., 1998; Garnett et al., 2012; Sato et al., 2005; Simões-Costa and Bronner, 2015; Simões-Costa et al., 2015; Steventon et al., 2009). In contrast, neural differentiation in the dorsal ectoderm results from the suppression of BMP, WNT, and Nodal signaling by secreted inhibitors including Noggin, Chordin, Follistatin, and DKK1 (del Barco Barrantes et al., 2003; Chambers et al., 2009; Fainsod et al., 1997; Li et al., 2013; Ozair et al., 2013; Sasai et al., 1995; Smith and Harland, 1992; Wilson et al., 1997).

Many studies have leveraged the knowledge gained from studies of model organisms to develop directed differentiation protocols using pluripotent human embryonic stem cells (hESCs). These protocols modulate the BMP and WNT signaling pathways to produce pure populations of desired cell fates within the ectoderm. As in *in vivo*, inhibition of TGFβ superfamily signaling is essential for differentiation to neural fates (Chambers et al., 2009), while appropriately timed introduction of BMP4 to the media can drive cells to epidermal and placodal fates (Dincer et al., 2013; Tchieu et al., 2017). Incorporating WNT or bMp4 has been shown to yield neural crest progenitors (Leung et al., 2016; Tchieu et al., 2017). Achieving pure populations of cells is likely essential in some applications, however the complexity of human development involves multiple self-organizing cell types whose patterning differs notably from other mammals and model organisms (Barry et al., 2017; Bock et al., 2018). Studying human ectodermal patterning requires developing *in vitro* models in which cells self-organize and spatiotemporal patterns emerge reproducibly in a fashion similar to that of the human embryo *in vivo*.

Recently, our lab and others have demonstrated that hESCs possess a remarkable ability to self-organize, and that this ability can be harnessed by confining them to micropatterned surfaces, so that embryonic fate patterns are generated *in vitro* (Etoc et al., 2016; Knight et al., 2018; Warmflash et al., 2014; Xue et al., 2018; Chhabra et al., 2018; Heemskerk et al., 2017). In this study, we developed a micropatterned platform in which cells self-organize to all four main fates within the ectoderm, and we use this system to dissect the spatiotemporal signaling events that pattern the human ectoderm. We demonstrate if hESCs are directed to adopt the ectodermal germ layer by inhibiting Nodal signaling, subsequent application of BMP4 to micropatterned colonies is sufficient to trigger dorsal-ventral patterning of multiple cell fates. Furthermore, we identified the duration of WNT signaling as a crucial control parameter that modulates the fate composition of ectodermal patterns. Finally, we used this system to understand how BMP and WNT signaling pattern the ectoderm. In particular, we show that cells are sensitive to the relative, rather than absolute, levels of BMP and WNT, and that this system can be used to dissect interactions between signaling and transcription factor networks in neural crest differentiation. We also show that taking the knowledge gained from this system into account allows us to improve on current protocols for placodal differentiation. Taken together, this study presents a novel self-organized ectodermal patterning system, and shows the power of this system for understanding human ectoderm development and improving differentiation towards desired ectodermal fates.

## Results

### Determining the window of competence for ectodermal patterning

We sought to create an in vitro model system to study the formation of patterned human ectodermal tissue. In analogy with our previous work creating gastrulation-stage patterns *in vitro*, we reasoned that geometrically confined hESCs treated with appropriately timed exogenous stimuli would create selforganized patterns of fates within the ectodermal germ layer. In both model organisms and embryonic stem cells, Nodal inhibition has been shown to be crucial for preventing mesendoderm differentiation, allowing cells to adopt ectodermal fates, while BMP signaling is responsible for generating the dorsalventral pattern within the ectoderm (Li et al., 2013, 2015; Liu et al., 2018). Thus, we hypothesized that following a two-stage protocol where hESCs are initially induced by Nodal inhibition and subsequently stimulated by BMP4 would cause them to exclusively adopt ectodermal fates. Although we expect these fates to be spatially disorganized in standard culture, this would provide a starting point for micropatterning experiments that test whether geometric confinement leads to ordered emergence of the same set of fates. Previous work in mouse and hESCs, has shown that prolonged Nodal inhibition leads to commitment to the neural fate (Li et al., 2015; Liu et al., 2018; Smith et al., 2008), so we specifically sought a temporal window in which cells are committed to the ectoderm but not exclusively to neural fates.

Compared to pluripotent cells, ectodermal progenitors are characterized by lower levels of pluripotency markers OCT4 and NANOG, high levels of SOX2, and the absence of neural specific genes such as PAX6 (Chambers et al., 2009; Li et al., 2015; Liu et al., 2018; Wang et al., 2012). To determine when hESCs enter this state, we examined these markers as a function of the duration of Nodal inhibition, achieved by growing cells in N2B27 media supplemented with 10 μm of SB431542, a small molecule ALK4/5/7 receptor-kinase inhibitor (hereafter called ectodermal induction media). By day 2, OCT4 and NANOG were fully repressed, while SOX2 expression levels remained comparable to those in the pluripotent state (Fig 1B,D, Supp1B). On day 3, we observed low and heterogeneous expression levels of PAX6 and NCAD (Fig 1A, Supp1A). By day 4, both PAX6 and NCAD expression increased in intensity and were homogeneously expressed (Fig 1A,B). Together, these data indicate that hESCs acquire a transient ectodermal progenitor state (SOX2+/NANOG-/OCT-/PAX6-/NCAD-) by day 2 of Nodal inhibition, but quickly differentiate towards the neural lineage upon continued Nodal inhibition.

**Figure 1:**
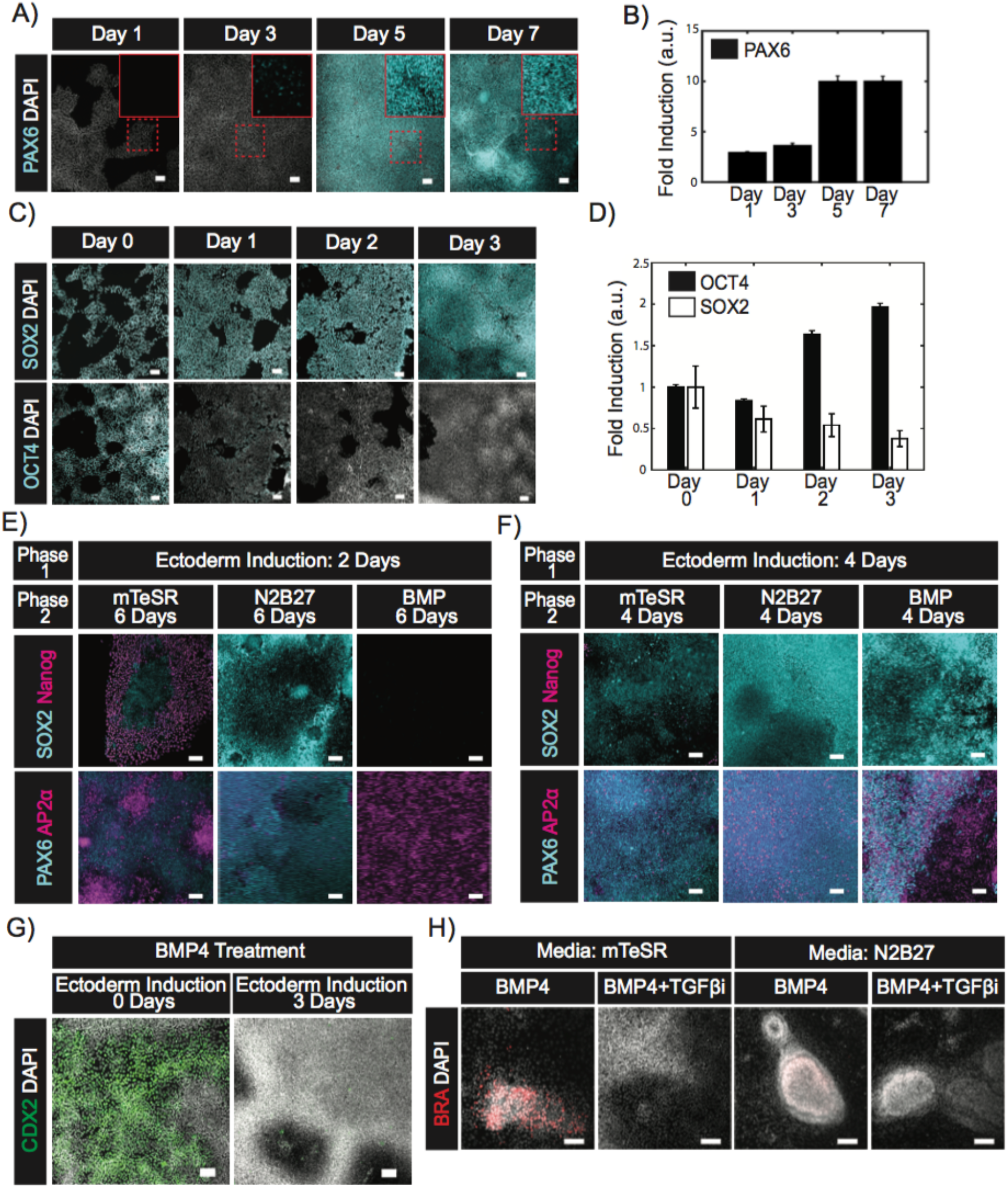
During neural differentiation cells transition through an ectodermal progenitor state before commitment. (A,C) Representative images of cells fixed on the indicated days during the course of Nodal inhibition and immunostained for the neural progenitor marker PAX6 (A) or the pluripotent markers SOX2 and OCT4 (C). Solid red inset in A is an enlarged image of the region marked by a dashed red box. (B,D) Quantification represents the fold change in average intensities of PAX6 (B) or SOX2 and OCT4 (D) normalized to DAPI for the indicated days relative to day 0. N=4 images. (E-F) Representative images of cells immunofluorescencently costained for AP2α/PAX6 or SOX2/NANOG at the conclusion of a two-phase induction protocol. Cells were initially differentiated in phase 1 for either 2 (E) or 4 (F) days in Nodal inhibition media and then treated with the indicated culture conditions for the subsequent 6 (E) or 4 (F) days in phase 2. (G) Representative images of cells immunoflourescently labeled for CDX2. All cells were treated with BMP4 for two days following either no prior treatment or 3 days of Nodal inhibition. (H) Representative images of cells immunoflourescently labeled for BRA at the conclusion of the experiment. All cells were initially induced for 3 days in Nodal inhibition media. Thereafter, the media was exchanged for treatment conditions indicated in the banner above the corresponding images. First row banners indicate the base media and the second banner row indicates the signals added to the base media. Scale bar = 100 μm

We next assessed the differentiation potential of hESCs as they transition from ectodermal to neural progenitors by exchanging the ectodermal induction media for a neutral (N2B27), pluripotency maintaining (mTeSR), or non-neural differentiation (N2B27+BMP4) media. Switching to a non-neural differentiation media on day 2 directed cells to non-neural ectoderm (AP2α+/PAX6-/SOX2-), while neutral media allowed most cells to adopt a neural fate (PAX6+/SOX2+/ AP2α-) (Fig 1E). Reverting to mTeSR on day 2 yielded a mixed population of fates consisting of neural, non-neural and pluripotent cells (Fig 1c). In contrast, exchanging ectodermal for non-neural induction media on day 4 generated clear territories of neural (PAX6+/AP2a-) and non-neural fates (PAX6-/AP2a+) by day 8, while media changed to neutral or mTeSR generated an expression profile consistent with the acquisition of a neural progenitor fate (high PAX6, low AP2α) (Fig 1F) (Chambers et al., 2009a). Collectively these data demonstrate that all hESCs remained competent to revert to pluripotency or differentiate towards nonneural fates after two days of ectodermal induction, while after 4 days all cells had committed to the ectodermal lineage and a fraction of cells had committed towards the neural fate.

Pluripotent cells give rise to mesodermal and extraembryonic fates in response to BMP4 (Bernardo et al., 2011; Nemashkalo et al., 2017; Xu et al., 2002). We next tested whether cells lose competence to differentiate to these fates following 3 days of ectodermal induction. Pluripotent hESCs treated with BMP4 for 2 days upregulated CDX2, however, its expression was absent if cells were first grown in ectoderm induction media for 3 days (Fig 1G). Additionally, we noted that following 3 days of ectodermal induction, 2 days of BMP4 treatment failed to up regulate mesodermal fates, defined by BRACHYURY (BRA) expression, when introduced with or without the Nodal inhibitor, SB, in N2B27 media. BRA+ cells were detected in the case when cells were treated with BMP4 in mTeSR media following three days of ectoderm induction. However, addition of SB to mTeSR blocked mesodermal differentiation (Fig 1H). This indicated that during the first three days of ectodermal differentiation, cells lose the potential to differentiate towards extraembryonic fates, but maintain competence to differentiate to mesodermal fates. Mesodermal differentiation requires an exogenous source of TGFβ and/or FGF, consistent with earlier reports on the requirements for mesoderm induction through BMP signaling in pluripotent stem cells (Bernardo et al., 2011; Yu et al., 2011)

### A two-step induction protocol generates patterns of human ectodermal fates in vitro

The previous experiments demonstrated that a properly timed two-step induction protocol with continuous Nodal inhibition could give rise to multiple ectodermal fates. We reasoned that carrying out this induction in geometrically confined colonies might lead to reproducible spatial patterning. We seeded hESCs on micropatterned surfaces and grew them for 72 hours in ectodermal induction media. Patterning was initiated with the addition of 50 ng/ml BMP4 in the presence of SB (10μm) and fixed following 72 hours of treatment. At the conclusion of the induction period, stem cell colonies routinely formed multilayered cellular structures with a dense ring of cells aggregating at a reproducible radius within the colony. We evaluated the expression of lineage specific markers within these micropatterned colonies.

Cells at the center of the micropatterned colonies differentiated to neural lineages (PAX6+/NCAD+/OTX2+/ECAD-/ISL1-) while those closer to the edge differentiated to non-neural ectoderm (PAX6-/NCAD-/OTX2-/ECAD+/ISL1+) (Fig 2A, Supp 2A) (Leung et al., 2013; Ozair et al., 2013; Tchieu et al., 2017). A ring of cells expressing the neural crest markers SOX9 and PAX3 was expressed between, and partially overlapped with the neural and non-neural domains (Fig 2a)(Betters et al., 2010; Bhatt et al., 2013; Monsoro-Burq et al., 2005; Spokony et al., 2002). Next to these cells was a ring of cells positive for the placodal marker SIX1 (Fig 2A, Supp 2B)(Dincer et al., 2013; Duncan and Fritzsch, 2013; Schlosser, 2006). These SIX1+ placodal cells were a subset of the ECAD+/ISL1 + population, and the SIX1-cells in this population represent epidermal precursors (Dincer et al., 2013; Tchieu et al., 2017).

**Figure 2:**
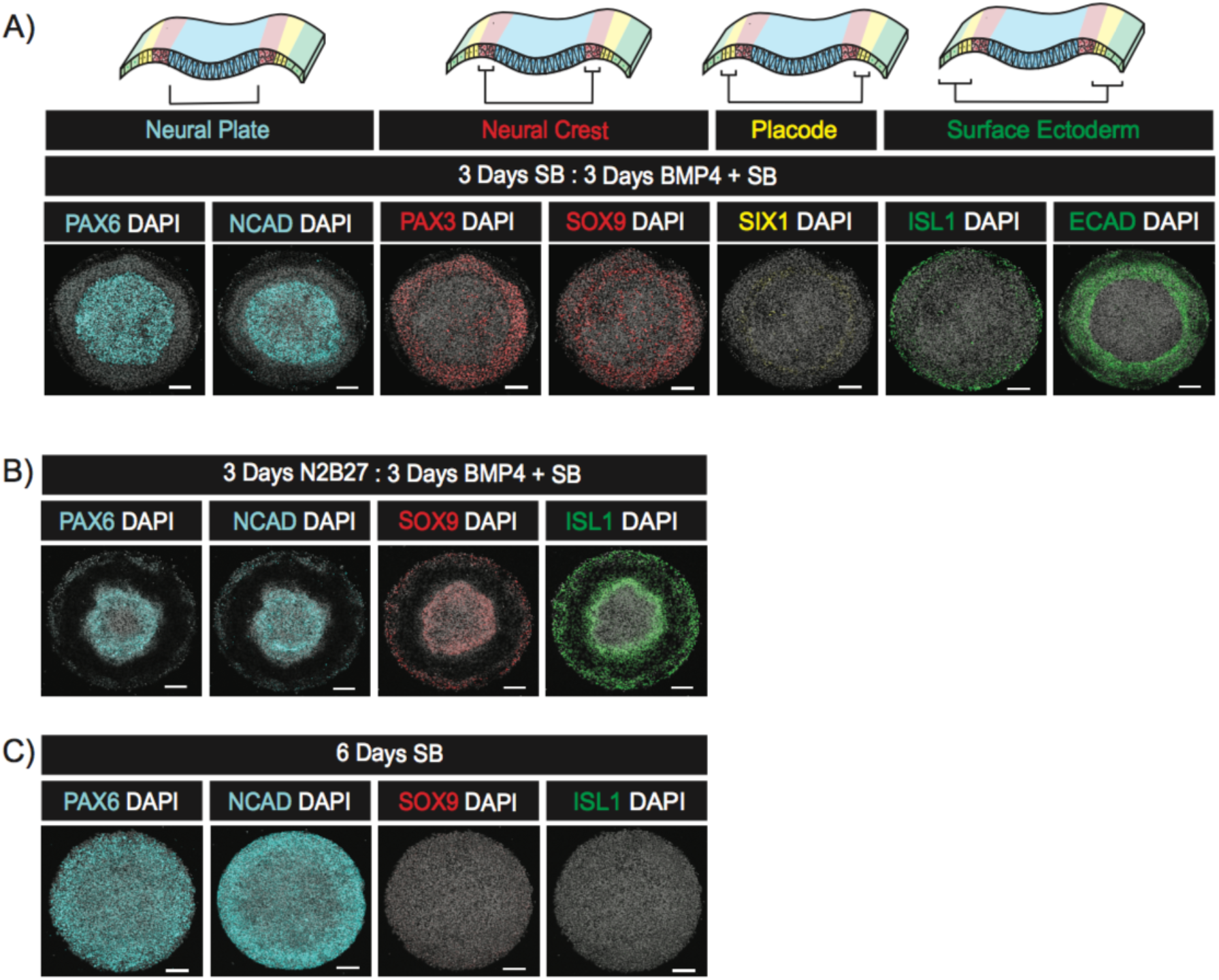
A two-phase protocol generates self-organized patterns of ectodermal fates. (A-C) Representative images of colonies stained with the indicated antibodies following treatment with either a two step protocol consisting of 3 days in ectoderm induction media and three days in N2B27+BMP+SB (A), the same protocol except the first three days were in N2B27 alone (B), or 6 days of ectoderm induction media (C). The top row in (A) shows a schematic of the organization of fates in the anterior ectoderm.

We next evaluated the necessity of Nodal inhibition in the first phase and BMP stimulation in the second to achieve ectoderm patterns. Removing SB from the first phase resulted in a contraction of the PAX6+ domain, the expansion of SOX9 expression to the colony edge, and the coexpression of SOX9 in the remaining PAX6+ cells at the colony center (Fig 2B). Thus, inhibition of Nodal signaling during the first three days is required to instruct future neural fates in the center of the colony upon BMP4 treatment.

Colonies that were grown for 6 days in ectoderm induction media without added BMP4 expressed PAX6 and NCAD throughout, while the surface ectoderm marker ISL1 was completely absent at the colony edge (Fig 2C). We observed a small number of SOX9 expressing cells near the colony edge but without a clear pattern. Thus, exogenous BMP4 stimulation is required for surface ectoderm differentiation and self-organized patterning.

### Duration of endogenous WNT signaling affects the composition of ectodermal tissue

While the protocol above yielded self-organized patterns consisting of four ectodermal lineages, the neural crest frequently formed in a broad domain without sharp boundaries, and there was limited placodal differentiation. Since WNT signaling is crucial for neural crest differentiation, we hypothesized that WNT ligands are endogenously produced and secreted, and that modulating these signals could alter patterning outcomes (Kurek et al., 2015). We therefore introduced IWP2, a small molecule inhibitor of WNT ligand secretion, at varying times during the two-step induction protocol (Chen et al., 2009). Introduction of IWP2 for all 6 days, or for the final 3 days, concurrently with BMP treatment, completely inhibited the expression of the neural crest markers PAX3 and SOX9 (Fig 3A, SuppFig 3A). Delaying IWP2 treatment for 12 hours following BMP4 addition was sufficient to initiate neural crest marker expression in a thin ring at the intersection of the neural and surface ectoderm (Fig 3A, SuppFig 3A). Allowing longer durations of WNT signaling by delaying the introduction of IWP2 caused an expansion of the neural crest territory from the edge of the neural domain inwards. Longer periods of endogenous WNT signaling also reduced the expression levels of the placodal marker SIX1 (Fig 3A). Together, our results demonstrate that WNT ligands are endogenously secreted and operate to alter the composition of fates within ectodermal tissue. Continuous inhibition of WNT secretion results in colonies consisting of domains of neural, placodal and future epidermal lineages, and lacking neural crest. WNT signaling creates the domain of neural crest cells, and longer durations of signaling expand this domain at the expense of neural and placodal fates.

**Figure 3:**
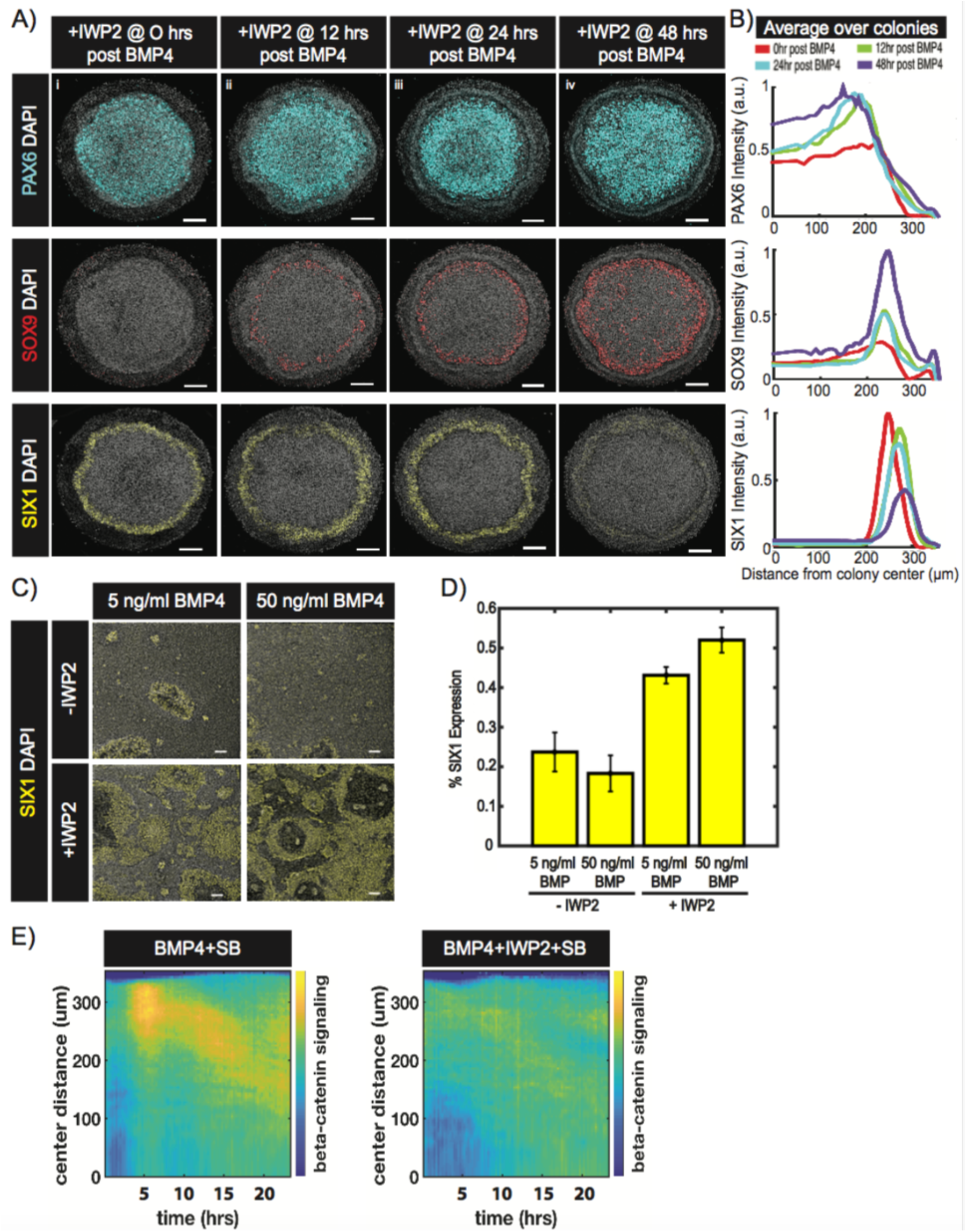
Endogenous WNT ligands drive differentiation to neural crest at the expense of placodal fates. (A) Representative images of colonies immunostained for PAX6, SOX9, or SIX1. Colonies were initially induced for 3 days in ectoderm induction media and then subsequently differentiated for 3 days in N2B27 media containing BMP4 and SB. The time between BMP4 and IWP2 addition is indicated in the banner above the corresponding image. Colony diameter = 700 μm.(B) Quantification of images in (a) represents average nuclear intensities of indicated markers normalized to DAPI as a function of radial position. N=3. (C) Representative images of cells in standard culture immunostained for SIX1. Cells were initially induced for 3 days in ectoderm induction media and then treated with either 5 or 50 ng/ml of BMP4 in media with (+) or without (-) IWP2 for the subsequent 3 days. Scalebar = 100 μm. (D) Quantitative representation of SIX1 expression normalized to the number of total cells in (c). N=4 images. (E) Kymograph of β-catenin signaling activity over a 24-hour period (on day 4) in micropatterned colonies initially treated in Nodal inhibition media for 3 days and then with the indicated signaling conditions. N=4 colonies

The above experiments demonstrated that delaying IWP2 addition for 12 hours after stimulation with BMP4 was sufficient to initiate neural crest differentiation. We reasoned that WNT signaling must begin prior to this time point to drive this differentiation. To test this, we monitored WNT pathway dynamics using a cell line with GFP fused to β-catenin at the endogenous locus. The onset of WNT signaling occurs within 10 hours of BMP4 treatment and its activity is strongly reduced by adding IWP2 simultaneously with BMP4 (Fig 3E, Supp Fig3C) (Massey et al., 2018). WNT activity was strongest in the region of future neural crest and moved inward towards the center of the colony in time. Thus, WNT signaling required neural crest differentiation is activated shortly after BMP4 addition.

### Wnt inhibition improves placode differentiation protocols

Placodes are unique cells of the ectoderm that give rise to the sensory organs. Previous reports have suggested that the specification of placodal fates by BMP is dose-dependent with high levels (> 10 ng/ml) inhibiting its expression and intermediate levels (~5 ng.ml) optimally inducing SIX1 (Leung et al., 2013; Tchieu et al., 2017). An alternative hypothesis is that higher levels of BMP induce greater WNT signaling which diverts potential placodal cells to neural crest. To distinguish between these hypotheses, hESCs were seeded in standard culture and initially induced for 3 days in ectoderm induction media. Thereafter, the cells were treated with 5 or 50 ng/ml of BMP4 either with or without IWP2 for 3 days. Treatments with IWP2 increased the fraction of SIX1+ cells at both low and high doses of BMP4 (Fig 3C,D). As previously reported, without IWP2, 5 ng/ml of BMP4 yielded a higher fraction of placodal cells than 50 ng/ml, although this difference was not statistically significant (p = 0.15) (Tchieu et al., 2017). In the presence of IWP2, this difference was abolished (Fig 3C,D). Thus, the inhibition of placodal differentiation at higher doses of BMP4 is caused by stronger activation of WNT signaling rather than resulting from BMP signaling directly. Preventing WNT secretion enables efficient placodal differentiation independently of the BMP4 dose.

### The width of the domain of placodal and epidermal progenitors is controlled by BMP4 concentration

Next we sought to better understand the correspondence between the level of exogenous BMP4 and patterning outcomes. In all cases, we implemented a three-step induction protocol with IWP2 added one day after BMP4 so that we achieved robust patterns of all four ectodermal fates at high BMP4 doses. At high concentrations (50 and 5 ng/ml), we observed the spatial organization of ectoderm progenitors described above (Fig 4A i,ii). Unexpectedly, we observed expression of AP2α within the neural domain, whereas *in vivo*, its expression pattern is limited to the future neural crest and surface ectodermal fates (Fig 4A i,ii) (Luo et al., 2002, 2003; Mitchell et al., 1991). With lower doses of BMP4, AP2α expression was restricted to the edge of the colony and was mutually exclusive with PAX6 expression. The width of the AP2α expressing territory depended on the BMP4 dose, with higher doses giving wider regions of expression (Fig 4A iii,iv,v). Consistent with this trend, lower doses of BMP4 lead to an expansion of the PAX6+/NCAD+ neural domain at the expense of the ISL1+/ECAD+ surface ectoderm (Fig 4A iii,iv,v,B,C). Additionally, coordinated with this expansion of the neural domain, we observed a shift of both neural crest and placodal progenitors towards the edge of the colony (Fig 4A,B,C). Lowering BMP4 concentration to 0.2 ng/ml led to patterns consisting only of neural in the center and neural crest at the edge of the colony. The loss of placodal fates (ISL1+/SIX1+) was not due to WNT signaling rather than insufficient BMP4 signaling, because inhibition of WNT secretion throughout the induction protocol did not rescue the expression of placodal (ISL1+/SIX1+) or surface ectoderm (ISL1+/SIX1-) fates (SuppFig 4A,B). Together, these data suggest there is a critical level of BMP4 that is necessary to initiate surface ectodermal differentiation, and that the width of the territory of surface ectoderm differentiation at the edge of the colony increases with BMP4 concentration.

**Figure 4:**
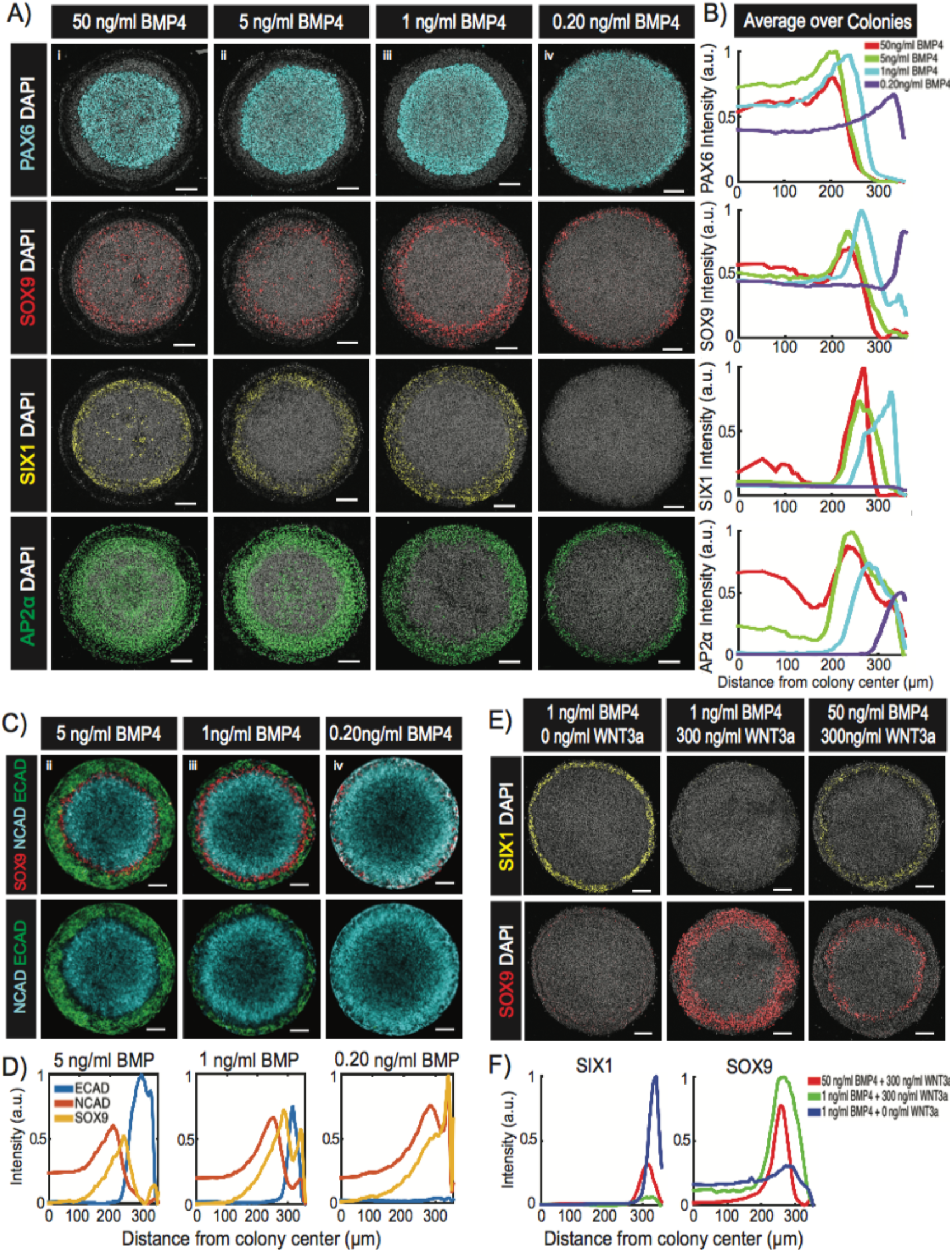
Cell fates are determined from the relative levels of BMP4 and WNT3a. (A,C) Representative images of colonies with the indicated single (A) or multiplexed (C) immunolabels following a three-step ectoderm induction protocol and patterned with (i) 50 ng/ml BMP4, (ii) 5 ng/ml BMP4, (iii) 1 ng/ml BMP4 or (iv) 0.20 ng/ml BMP4 in N2B27 media with SB. Colony diameter = 700 μm. (B,D) Quantifications of images in A (B) and C (D) represents the average nuclear intensities of indicated markers normalized to DAPI as a function of radial position. N= 3 colonies. (E) Representative images of colonies immunostained for SOX9 or SIX1. Cells were initially differentiated in ectoderm induction media for 3 days and subsequently induced in media containing IWP2 with the indicated levels of BMP4 and WNT3a in the overhead banner. Colony diameter= 700 μm. Scalebar = 100 μm. (F) Quantification of images in E represent the intensities of the indicated markers normalized to DAPI as a function of distance from the colony center. N=3 colonies.

### The relative levels of BMP and WNT instruct the decision between neural crest and placode

The results above argue that cells integrate information from the BMP and WNT pathways to create patterns within the ectoderm. We hypothesized that at the edge of the colony, a region competent to form both neural crest and placodes (Fig 4A iii,iv), the decision between these fates depends on the relative, rather than absolute, levels of WNT and BMP signals. To test this, micropatterened hESC colonies were stimulated with different relative concentrations of exogenous BMP4 and WNT3a in the second phase of the induction protocol. To avoid endogenous WNT signaling complicating the interpretation of these experiments, IWP2 was used throughout the second phase in all treatments. SIX1 is induced at the edge of the colony by 1 ng/ml of BMP4, and was significantly suppressed when 300 ng/ml of WNT3a and 1 ng/ml of BMP4 were added together with cells at the border instead adopting SOX9+ neural crest fates (Fig 4E). Raising the BMP4 concentration to 50 ng/ml while maintaining 300 ng/ml of WNT3a rescued the expression of SIX1 at the edge of the micropattern and reduced that of SOX9 (Fig 4E). These data show that cell-fate decisions within the ectoderm are determined by the relative concentrations of BMP and WNT ligands.

### Dynamics of cell-fate decisions in micropatterned ectoderm

We next studied the spatiotemporal dynamics of key markers associated with fate acquisition during the course of ectodermal patterning. hESCs were seeded on a micropatterned surface, differentiated using a three-step induction protocol with 1 ng/ml of BMP4 added on day 4 and IWP2 added one day later. Cells were fixed on days 5, 6 and 7 and the expression of fate markers evaluated. We found that the expression of SOX2, initially an ectoderm marker and later a neuroectoderm marker, was expressed throughout the colony on day 4, however, on days 6 and 7, high SOX2 levels were progressively restricted to the center of the colony while the edges maintained a lower level of expression (Fig 5) (Li et al., 2015; Papanayotou et al., 2008; Zhang et al., 2010). NCAD and PAX6 expression were initiated in the center of the colony on day 5 and maintained in this domain throughout the induction protocol (Fig 5). The non-neural marker, AP2α was found in a ring at the colony border throughout the induction period, whereas ISL1, a marker of surface ectoderm, was detected later on day 6 and maintained on day 7 in the same region (Fig 5). ECAD, a marker found in both pluripotent and surface ectodermal cells, was weakly expressed in the center of the colony while higher levels were observed at the colony edge on day 5. Thereafter, ECAD expression was lost in the center and maintained at the colony edge on days 6 and 7 (Fig 5). For placodal markers, we found GATA3 expression at the colony edge on days 5 and 6, but its levels were reduced by day 7, recapitulating the transient GATA3 expression observed during placode development in model organisms (Fig 5) (Bhat et al., 2013; Groves and LaBonne, 2014; Schlosser, 2006). The expression of SIX1 was found at the colony edge on day 6, 24 hours later than GATA3, and was maintained on day 7 (Fig 5). The neural crest markers SOX9 and PAX3 were detected at low levels on day 5 and increased over time with the expression domain remaining in the same radial position (Fig 5). Thus most markers initially appear in their final domain of expression, and expression levels rise and cell fate boundaries are progressively sharpened in time.

**Figure 5:**
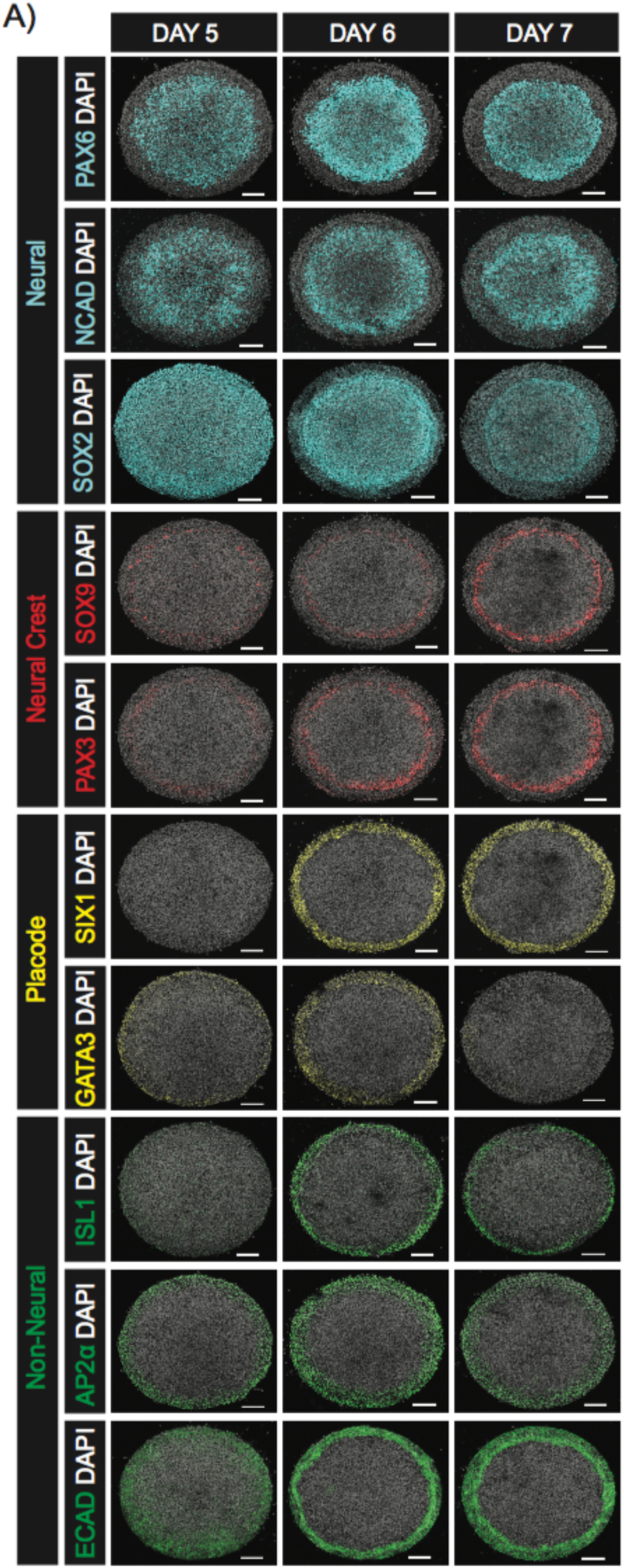
Spatiotemporal characterization of human ectoderm pattern emergence. (A) Representative images of colonies fixed on the indicated days and immunostained with the indicated antibodies. Cells were differentiated using a three-step ectoderm induction protocol and patterned with 1 ng/ml BMP4. Colony diameter = 700 μm. Scalebar = 100 μm

### Signaling history influences the interpretation of BMP4 by hESCs

During the first three days of induction, a small molecule Nodal inhibitor is applied without stimulating other pathways, however, endogenous signals may be required during this time. We first assessed BMP signaling through C-terminally phosphorylated SMAD1/5/8 levels (pSmad1/5/8), a proximal read out of BMP activity (Hoodless et al., 1996; Kawai et al., 2000; Nishimura et al., 1998). We observed a prepattern with active endogenous BMP signaling at the colony edge. This prepattern was specific to BMP signaling as it was abolished by treatment with the BMP inhibitor LDN (Fig 6A) (Yu et al., 2008). To assess the functional significance of this prepattern, cells were induced for three days with SB+LDN and then treated with BMP4. Under these conditions, colonies created patterns consisting only of neural crest at the colony edge and neural in the center (Fig 6B). However, both neural and surface ectodermal fates were observed if colonies were pretreated in SB+LDN for only 1 or 2 days prior to BMP stimulation (Fig 6b) with the width of the surface ectoderm domain reduced the longer BMP signaling was inhibited. Thus, a prepattern of endogenous BMP signaling is required to preserve competence for surface ectodermal and placodal differentiation.

**Figure 6:**
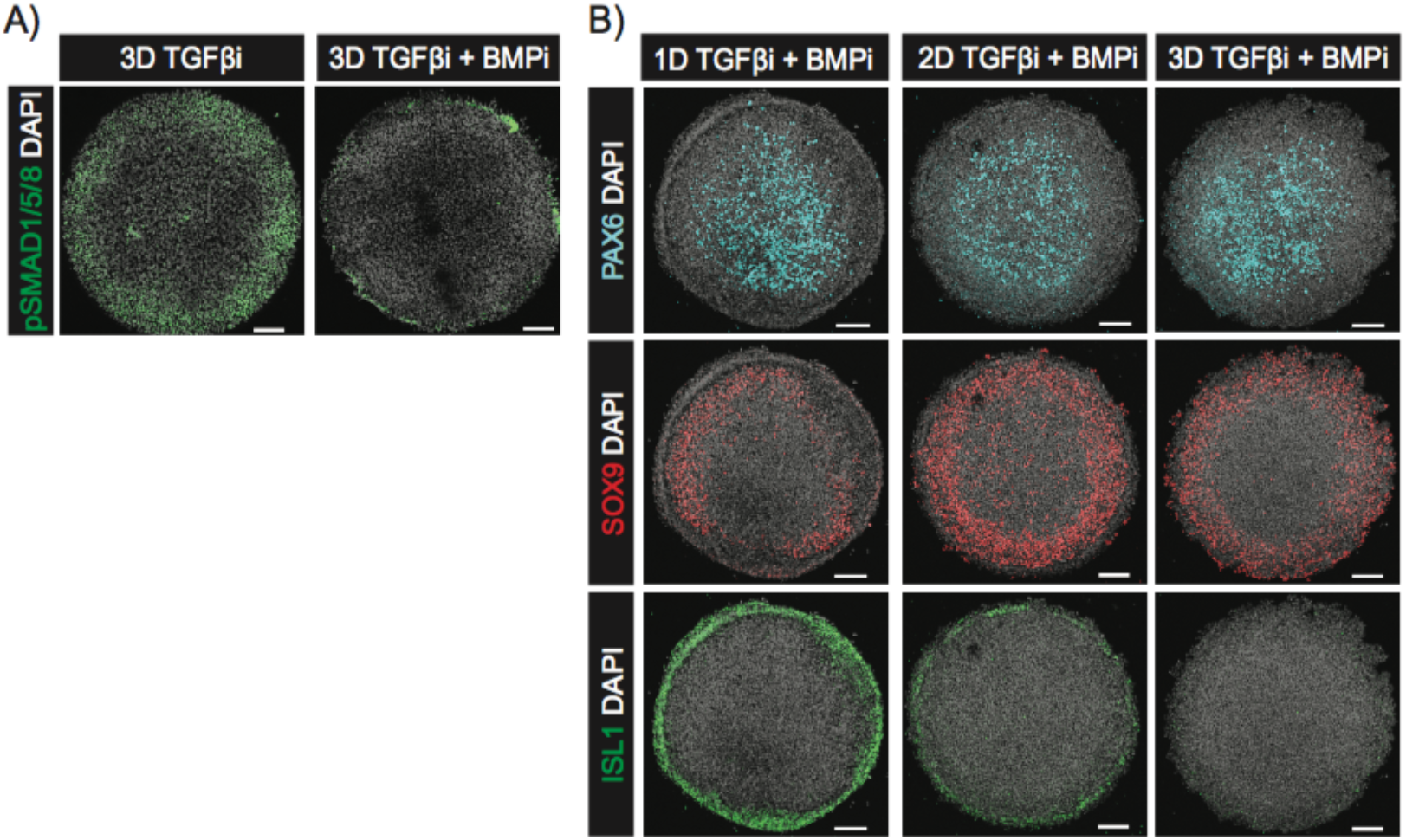
Endogenous BMP signaling prior to BMP4 treatment is required for surface ectodermal differentiation. (A-B) Representative images of colonies immunostained for the indicated antibodies on day 3 following induction in media containing SB only or in combination with LDN (A), and on day 6 following 1, 2 or 3 days of SB+LDN treatment in the first phase of induction prior to BMP4 treatment for 3 days. Conditions that removed duel inhibition before day 3 had media replaced with N2B27 + SB such that all conditions were in culture for the same duration prior to the introduction of BMP4 (B). Colony diameter = 800 μm (A); 700 μm (B). Scalebar = 100 μm.

### Combinatorial signaling logic guiding neural crest specification

Experimental evidence from model organisms suggests that the transcription factors SOX9, PAX3 and AP2α are required for the specification of neural crest (Betters et al., 2010; Bhatt et al., 2013; Garnett et al., 2012; Leung et al., 2016; Monsoro-Burq et al., 2005; Sato et al., 2005; Simões-Costa and Bronner, 2015). The regulatory links between these genes and others have been assembled into gene regulatory networks, but it remains unclear how these networks combinatorially interpret signaling from the BMP and WNT pathways. We aimed to leverage our *in vitro* model to dissect the contributions from BMP and WNT signaling pathways to the specification of neural crest fates. To accomplish this, cells were first induced with either SB alone or in combination with LDN for 3 days and then BMP4 or WNT3a was added alone or in combination with an inhibitor to the second pathway (Fig 7A,B). Colonies treated with WNT3a (300 ng/ml) following 3 days of Nodal inhibition showed overlapping expression of SOX9, PAX3 and AP2α at the colony edge (Fig 7A ii). However, inhibition of endogenous BMP (Fig 6A) prior to WNT stimulation leads to the expression of PAX3 without AP2α or SOX9 at the colony edge (Fig 7A i, 7B i). Conversely, inhibiting WNT at the time of BMP treatment resulted in the upregulation of AP2α but not PAX3 or SOX9 (Fig 7A iii). By isolating the activity of each pathway, our data suggest that BMP signaling contributes to the expression of AP2α, while WNT signaling induces the expression of PAX3. Both signals are required for expression of the mature neural crest marker SOX9.

**Figure 7:**
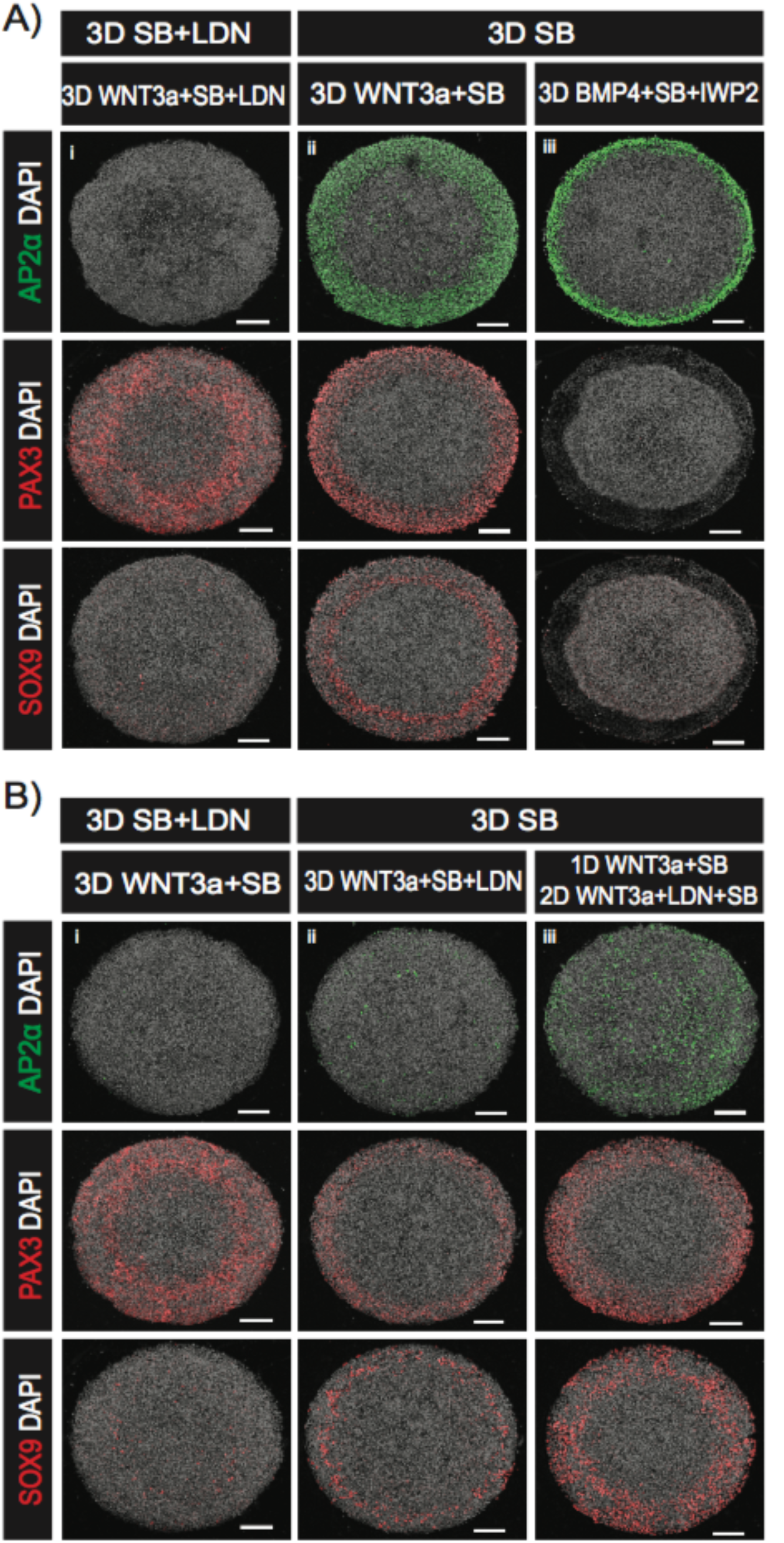
Dissecting the logic connecting BMP and WNT signaling to the transcription factors AP2a, PAX3, and SOX9. (A-B) Representative images of colonies immunostained for AP2α, PAX3 or SOX9. The top banner indicates the signaling conditions for the first three days (3D) while the banner in the second row indicates the signaling conditions for the subsequent three days. (i) 3 days SB: 3 days WNT3a+SB, (ii) 3 days SB: 1 day WNT3a+SB: 2 days WNT3a+LDN+SB, (iii) 3 days SB: 3 days WNT3a+SB+LDN (A). (i) 3 days SB+LDN: 3 days WNT3a+SB, (ii) 3 days SB: 3 days WNT3a+SB+LDN, (iii) 3 days SB: 1 day WNT3a+SB: 2 days WNT3a+LDN+SB (B). Colony diameter = 700 μm. Scalebar = 100 μm.

Lastly, we evaluated the temporal requirement for endogenous BMP signaling in generating neural crest at the colony borders in response to WNT stimulation (Fig 7B ii,iii). Introduction of the BMP inhibitor LDN concurrently or 24 hours post WNT3a stimulation reduced the expression of AP2α while maintaining that of PAX3 and SOX9 (Fig 7B ii,iii). Interestingly, the strength of AP2α reduction was dependent on the duration of BMP inhibition. These results suggest a previously unappreciated temporal requirement to specify neural crest fates: continuous endogenous BMP signaling is necessary to maintain AP2α expression, however, only a short duration in BMP signaling is required to initiate and maintain SOX9 once WNT is present. In the future, it will be interesting to determine whether the cells that are SOX9+PAX3+AP2α-are viable for further differentiation.

## Discussion

In this study, we developed a system in which hESCs create self-organized patterns of human neural, neural crest, placodal and epidermal progenitors on micropatterned surfaces with the same organization as found *in vivo* within the ectodermal germ layer along the dorsal-ventral axis. Cells were first specified to ectoderm through an initial phase of TGFβ inhibition, followed by a second phase in which patterning was initiated with BMP4, and third phase in which a WNT inhibitor was combined with BMP4 stimulation to prevent widespread activation of WNT. We leveraged our *in vitro* system to dissect the signaling interactions that underlie patterning and to improve current directed differentiation protocols for placodal cells. We showed that increasing WNT signaling duration leads to the expansion of neural crest at the expense of placodal and neural fates, and that cells sense the relative amounts of WNT and BMP signaling in choosing between placodal and neural crest lineages. Finally, we used the *in vitro* system to dissect the links between WNT and BMP signaling and the members of the gene regulatory network governing neural crest specification, links that have been difficult to untangle in model systems *in vivo* (Martik et al., 2017).

Recently, it was shown that hESCs grown on micropatterns in neural differentiation media would self-organize into two territories with neural cells in the center and neural crest in the periphery. The induction of neural crest at the colony edge relied on endogenous BMP signaling, and the authors suggested that BMP signaling is activated by mechanical effects at the colony edge (Xue et al., 2018). Micropatterns have also been used to study patterning within the neural cells themselves. During neural differentiation, cells organize themselves into rosette-like structures reminiscent of the neural tube (Elkabetz et al., 2008). This system provides an interesting opportunity to study the self-organizing properties of the neuro-epithelium, however, rosettes typically form in a variety of sizes in a disorganized fashion. A protocol was recently developed in which micropatterns are used so that a single rosette forms on each pattern, and the size of the patterns was shown to be a crucial control parameter (Knight et al., 2018). In the future, it will be interesting to investigate whether the center of our ectodermal micropatterns develops a rosette morphology with further differentiation, and whether self-organized mechanical cues play a role, especially since the WNT pathway plays a crucial role in our patterns and is a known mechanosensitive pathway (Benham-Pyle et al., 2015).

Although measurements of pSmad1/5/8 have proven difficult in mammalian embryos, experiments in Zebrafish and Xenopus have revealed a dorsal ventral gradient in signaling activity (Schohl and Fagotto, 2002; Tucker et al., 2008). In Xenopus, experiments with dissociated animal cap cells have suggested that the concentration of BMP can elicit multiple fates in a dose-dependent manner (Wilson et al., 1997). While here we observe a similar pattern of pSMAD1/5/8 along the radius of the micropatterned colonies, our results do not support a model in which cell fates along the dorsal ventral axis are patterned exclusively by the BMP dose (Nemashkalo et al., 2017; Chhabra et al., 2018). A minimum BMP4 concentration is necessary to initiate surface ectodermal differentiation, however, the decision to form neural crest or placodes requires the integration of BMP and WNT signals.

It is instructive to compare our results here with our previous work on selforganized germ layer differentiation in micropatterns. In both cases, patterning is initiated by an applied BMP4 signal, which combines with endogenous WNT signals. The edge fates (extraembryonic for gastrulation micropatterns, surface ectoderm in this case) are upregulated by BMP signaling while spatially intermediate fates (mesendoderm in the case of gastrulation, neural crest here), require the WNT signal. The fates at the center (ectoderm or neural) result from the inhibition of both signals. The essential difference in protocols is that the gastrulation patterns are generated directly from pluripotent hESCs while ectodermal patterns require a three-day induction towards ectoderm before application of BMP4. The similarity in these signaling response between the two cases suggests the possibility that a core signaling network comprising the BMP and WNT pathways is responsible for both patterns while the competence of the cells shifts in time. This allows the same BMP-WNT signaling circuit to elicit different fates during the patterning of germ layers at gastrulation and the patterning of the ectoderm slightly later in development. A crucial difference, however, is the role of Nodal signaling. While Nodal is essential for mesendoderm differentiation, it is inhibited throughout here. Given that cells are still capable of forming BRA-positive mesoderm after three days of ectodermal induction (Figure 1H), it would be interesting to determine whether removing Nodal inhibition or exogenously adding Nodal ligands would convert some of the ectodermal cells to mesodermal fates, and whether cell would form patterns of ectoderm and mesoderm under these conditions.

We showed that cells do not respond to a particular level of BMP or WNT signaling, but instead determine fate as a function of these pathways combinatorially. While elevating WNT signaling drives neural crest differentiation at the expense of placodes, the same levels of WNT are also compatible with placodal differentiation when BMP levels are also elevated. What is the role of this combinatorial control of cell fate? One possibility is that it is a means to allow diversification of cell fate by multiple pathways. This may be a more robust strategy than having cells read multiple levels off a single gradient. For example, rather than creating the four fates comprising the DV axis within the ectoderm by reading the precise levels of BMP4, cells may create the same four states by reading two pathways in a binary way. In agreement with this view, experimental and theoretical studies have questioned whether a single morphogen gradient or signaling pathway contains enough information to specify multiple fates (Gregor et al., 2007; Cheong et al., 2011). Additional sources of information may also be present in the signaling dynamics (Selimkhanov et al., 2014). More fine-grained mapping of the fate patterns as a function of signaling levels, as well as careful quantification of the signaling dynamics in each case will be required to fully understand this connection between signaling and cell fate.

The WNT pathway has been shown to play a critical role in anterior-posterior axis formation in the gastrulating embryo (Kiecker et al., 2001; Yamaguchi, 2001; Arnold et al., 2009). High WNT signaling is essential for primitive streak formation on the posterior side of the embryo, while WNT is inhibited anteriorly by secreted inhibitors from the anterior visceral endoderm (Kiecker et al., 2001; Yamaguchi, 2001; Arnold et al., 2009). Later, ongoing WNT signaling is required in the tailbud, and in vitro studies have suggested that longer Wnt exposure causes hESCs to adopt progressively more posterior fates (Knight et al., 2018). The shift in fate patterns that we observe when the duration of WNT signaling is modulated is consistent with this duration setting the position along the AP axis. When WNT signaling was inhibited throughout, the cell fate patterns resemble anterior ectoderm, and are composed of neural, placodal and future epidermal, but not neural crest, lineages. Permitting WNT activity generated ectodermal patterns consistent with the mid-brain, which are comprised of all four of these lineages. As the principle markers we use here are for fates along the DV axis, a more definitive assignment of AP position could be made by examining specific markers of AP position, such as OTX2, GBX2, and HOX genes. Recent studies support the idea that the posterior fates in the spinal cord are specified by a separate route from the anterior ones (Kiecker et al., 2001; Yamaguchi, 2001; Arnold et al., 2009; Gouti et al., 2014; Garriock et al., 2015; Lippmann et al., 2015) even prior to germ layer specification, and it would be informative to determine protocols for generating ectodermal patterns that mimic the posterior of the embryo.

A large body of work has discovered numerous transcription factors that are involved in different stages of neural plate border development and neural crest differentiation. For example, knockdown experiments in model organisms have demonstrated a requirement for PAX3 and AP2α in the specification of neural crest fates as defined by SOX9/10 expression (Bae et al., 2014; Luo et al., 2003). It is believed that both the BMP and WNT signaling pathways are required for the maturation of neural crest, but it has been difficult to unravel how these pathways are embedded in the gene regulatory network that specifies the neural crest *in vivo*. We showed that the micropatterned system is an attractive platform for dissecting these interactions due to the ease of manipulation and the removal of confounding factors resulting from effects on earlier development or on other tissues. Specifically, we showed that some individual genes require only BMP signaling, only Wnt signaling, or both, and that markers of more mature neural crest fall into this last category. We also noted that as long as endogenous BMP signaling is allowed during the first phase of the protocol, WNT activation alone is sufficient to activate all of these genes, so the requirement for BMP signaling in neural crest differentiation is only transient, while that for WNT is ongoing. This approach can be extended to gain a fuller picture of how signaling pathways direct a complex network of transcription factors in the specification and function of the neural crest.

This study has developed a novel system in which human embryonic stem cells generate the DV pattern within the ectoderm. This system can be used to gain an understanding of human ectodermal development and to improve directed differentiation protocols. Future studies can take advantage of this system together with live cell reporters of signaling and fate to understand in detail how ectodermal patterns are self-organized. Finally, our study provides proof-of-principle for extending the micropatterning approach beyond germ layer specification to later developmental events. As long as cells can be directed to adopt a progenitor cell fate, providing a reproducible geometric and chemical environment leads to patterning, and the patterns can then be tuned to yield a particular outcome. A similar approach can be used to generate patterns corresponding to endoderm or mesoderm, or to particular organs.

## STAR★Methods

### Contact for Reagent and Resource Sharing

Enquiries on reagents and resources should be directed to, and will be fulfilled by Aryeh Warmflash

### Experimental Model

#### Cell lines

Except for live cell imaging, all experiments were performed using ESI017 (obtained from ESI BIO, RRID: CVCL_B854, XX) human embryonic stem cell (hESC) line. For live imaging of Wnt signaling dynamics, the eSi017 GFP:β-catenin cell line as described in (Massey *et al.*, 2018) was used.

### Method details

#### Routine cell culture

All cells were grown in the chemically defined medium mTeSR1 in tissue culture dishes and kept at 37°C, 5% CO_2_ as described in (Nemashkalo *et al.*, 2017).

Cells were routinely passaged and checked for mycoplasma contamination also as described in (Nemashkalo *et al.*, 2017). In all experiments, cell passage number did not extend beyond 55.

#### Differentiation

*In standard culture*. ESI017 hESC cultures were washed two times with 1X PBS and disaggregated using accutase for 7 minutes at 37°C. Cells were subsequently harvested, centrifuged at 1000 rpm for 4 minutes and seeded on Matrigel coated Ibidi dishes at a density of 18,000 cells/cm^2^ in the presence of mTeSR media and ROCK inhibitor (ROCKi) (10μM). On the following day ROCKi was withdrawn from the media, and hESCs were allowed to expand in culture for 2 days in mTeSR. All inductions were initiated on the third day in N2B27 medium, which we prepared by filtering a mixture of 250 ml DMEM/12, 250 ml Neurobasal media, 2.5 ml N2 supplement, 5 ml B27 supplement without vitamin A, 5 ml glutamax, and 0.5 ml β-mercaptoethanol. N2B27 was supplement with growth factors and small molecules as indicated. In all experiments media was replenished daily.

*On micropatterns*. For micropatterned cell culture experiments, micropatterned glass CYTOOchips or 96-well CYTOOplates were coated with 5 μg/ml laminin-521 in 1X PBS with calcium and magnesium (PBS++) for 2.5 hours at 37°C. The laminin was then removed by first using serial dilutions without allowing the chip or well to dry (dilution 1:4 in PBS++, 5x), and then using one complete wash with PBS++. The chip or plate was either used immediately or stored for up to two weeks at 4°C with PBS++ covering the micropatterned surfaces.

Seeding of hESCs onto micropatterned surfaces was performed as follows. Stem cell cultures were washed two times with 1x PBS and disaggregated using accurate for 7 minutes at 37°C. Cells were subsequently harvested, centrifuged at 1000 rpm for 4 minutes and resuspended in mTeSR containing ROCKi. Cells were then seeded to laminin coated CYTOOchips (1 × 10^6^ cells in 2 ml of media) or laminin coated CYTOOplates (100,000 cell/well with 100 μl media/well), and then placed in the incubator for 45 minutes at 37°C. Cells were then washed twice with 1X PBS to remove cells bound nonspecifically to uncoated regions of the well or chip. To promote better adhesion to the micropatterned surfaces, all ectoderm inductions were initiated in the presence of mTeSR supplemented with SB431542. On the following day, mTeSR was withdrawn as the base media and replaced with N2B27 for the remainder of the experiment. Stem cell colonies were then further treated with reagents as described in the text. In all micropatterned experiments, the media was replenished daily with the described signaling conditions and fixed at the indicated times within the text.

#### Immunostaining

Immunostaining followed standard protocols as described in (Nemashkalo *et al.*, 2017). Primary and secondary antibodies were diluted in the blocking solution as described in (Warmflash et al., 2014; Nemashkalo et al., 2017). Dilutions are listed in the reagents table.

#### Imaging

##### Live cell imaging

GFP:β-catenin hESCs were seeded in multiple wells of a 96-well CYTOOplate and induced towards the ectoderm as described in the text above. Four colonies (700 μm) per condition were imaged with a 20X, NA 0.75 objective on a Olympus/Andor spinning disk confocal microscope. 5 z-sections were acquired per position every 30 minutes from 24 hours pre to 48 hours post BMP4 treatment. To remain consistent with fix cell experiments, the imaging was halted for a brief period (~1 hour) every 24 hours to replenish the media. In all cases, cells were maintained at 37°C and 5% CO_2_ throughout the duration of imaging.

##### Fixed cell imaging

All immunostaining data was acquired by imaging multiple z-sections at 10X (NA 0.40) from fixed cell colonies on CYTOOchips or CYTOOplates with an Olympus FV1200 laser scanning confocal microscope.

### Quantification and Analyses

All experiments were performed at least twice with consistent results. The data and analyses in each figure belong to one experiment. Sample size was not predetermined and no statistical tests were used to determine significance of results.

Circular colonies with a non-radial cell density pattern at the end of treatment were excluded from analyses. The number of colonies included in each analysis (N) is mentioned in the figure legends. For images taken at 10X magnification with multiple z-slices; background subtraction, maximum z projection and alignment were performed as described in (Warmflash *et al.*, 2014). The average intensity of a marker was calculated for each cell as the average of the immunofluorescent intensity in that cell normalized to the intensity of DAPI in the same cell. In cases in which the cells were too dense to effectively perform cell segmentation, averages were performed over nuclear pixels rather than cells.

### Data and Software

All the raw data is available upon request. MATLAB scripts for analyzing experimental data can be obtained from https://github.com/gbri140.

### Key Resources Table

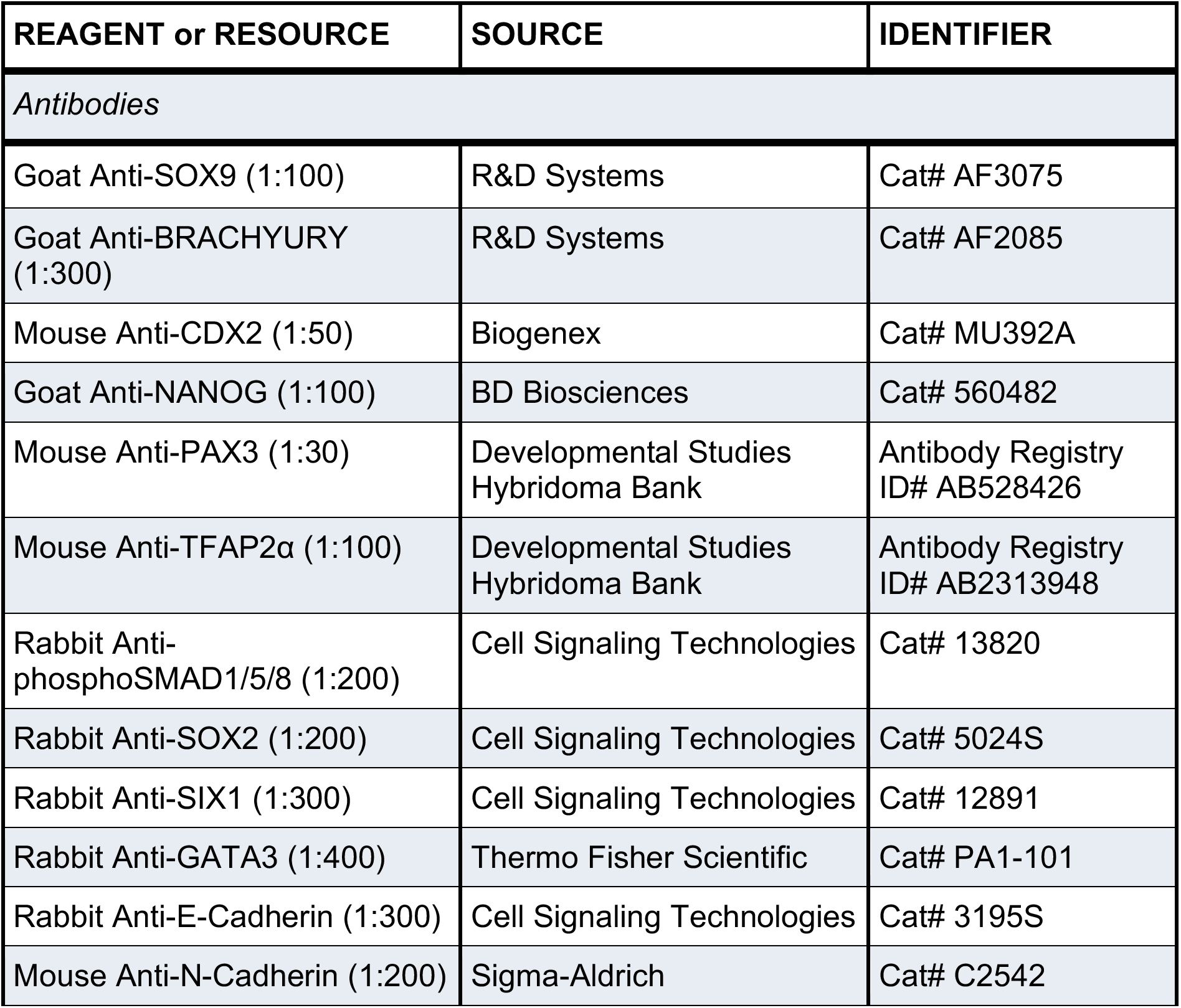
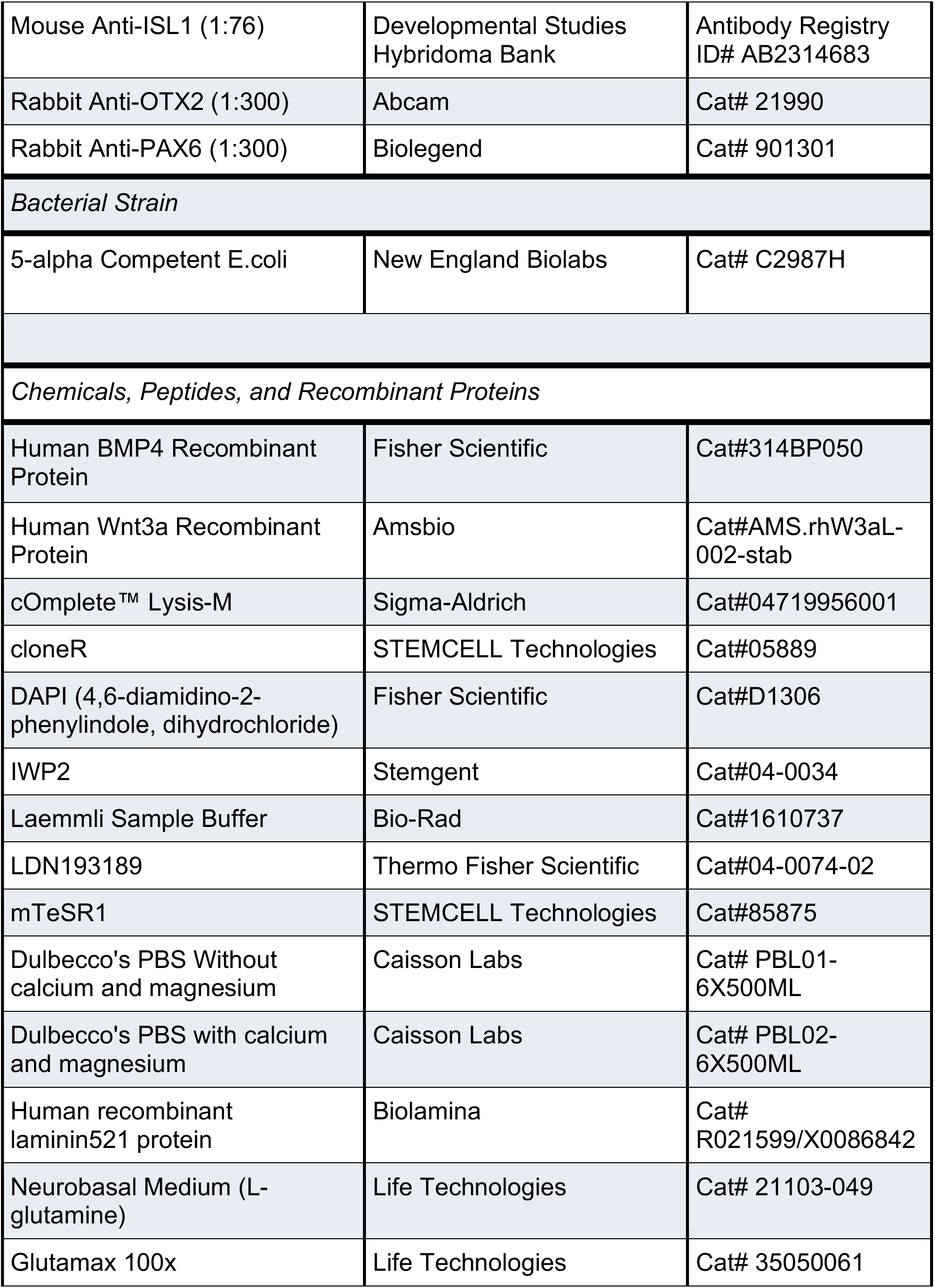
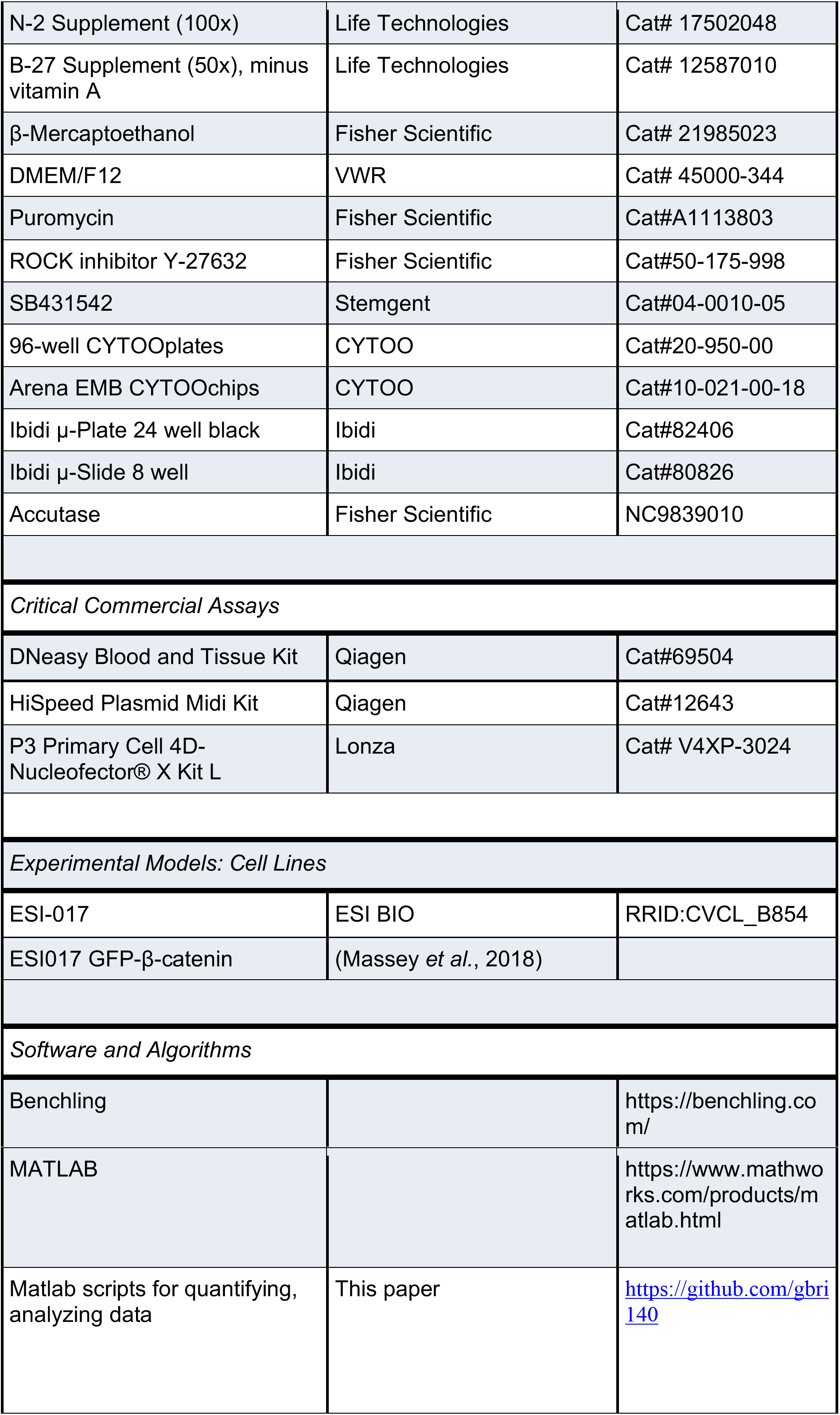

| REAGENT or RESOURCE | SOURCE | IDENTIFIER |
| --- | --- | --- |
| *Antibodies* |  |  |
| Goat Anti-SOX9 (1:100) | R&D Systems | Cat# AF3075 |
| Goat Anti-BRACHYURY (1:300) | R&D Systems | Cat# AF2085 |
| Mouse Anti-CDX2 (1:50) | Biogenex | Cat# MU392A |
| Goat Anti-NANOG (1:100) | BD Biosciences | Cat# 560482 |
| Mouse Anti-PAX3 (1:30) | Developmental Studies Hybridoma Bank | Antibody Registry ID# AB528426 |
| Mouse Anti-TFAP2α (1:100) | Developmental Studies Hybridoma Bank | Antibody Registry ID# AB2313948 |
| Rabbit Anti-phosphoSMAD1/5/8 (1:200) | Cell Signaling Technologies | Cat# 13820 |
| Rabbit Anti-SOX2 (1:200) | Cell Signaling Technologies | Cat# 5024S |
| Rabbit Anti-SIX1 (1:300) | Cell Signaling Technologies | Cat# 12891 |
| Rabbit Anti-GATA3 (1:400) | Thermo Fisher Scientific | Cat# PA1-101 |
| Rabbit Anti-E-Cadherin (1:300) | Cell Signaling Technologies | Cat# 3195S |
| Mouse Anti-N-Cadherin (1:200) | Sigma-Aldrich | Cat# C2542 |
| Mouse Anti-ISL1 (1:76) | Developmental Studies Hybridoma Bank | Antibody Registry ID# AB2314683 |
| Rabbit Anti-OTX2 (1:300) | Abcam | Cat# 21990 |
| Rabbit Anti-PAX6 (1:300) | Biolegend | Cat# 901301 |
| *Bacterial Strain* |  |  |
| 5-alpha Competent E.coli | New England Biolabs | Cat# C2987H |
| *Chemicals, Peptides, and Recombinant Proteins* |  |  |
| Human BMP4 Recombinant Protein | Fisher Scientific | Cat#314BP050 |
| Human Wnt3a Recombinant Protein | Amsbio | Cat#AMS.rhW3aL-002-stab |
| cOmplete™ Lysis-M | Sigma-Aldrich | Cat#04719956001 |
| cloneR | STEMCELL Technologies | Cat#05889 |
| DAPI (4,6-diamidino-2-phenylindole, dihydrochloride) | Fisher Scientific | Cat#D1306 |
| IWP2 | Stemgent | Cat#04-0034 |
| Laemmli Sample Buffer | Bio-Rad | Cat#1610737 |
| LDN193189 | Thermo Fisher Scientific | Cat#04-0074-02 |
| mTeSR1 | STEMCELL Technologies | Cat#85875 |
| Dulbecco's PBS Without calcium and magnesium | Caisson Labs | Cat# PBL01-6X500ML |
| Dulbecco's PBS with calcium and magnesium | Caisson Labs | Cat# PBL02-6X500ML |
| Human recombinant laminin521 protein | Biolamina | Cat# R021599/X0086842 |
| Neurobasal Medium (L-glutamine) | Life Technologies | Cat# 21103-049 |
| Glutamax 100x | Life Technologies | Cat# 35050061 |
| N-2 Supplement (100x) | Life Technologies | Cat# 17502048 |
| B-27 Supplement (50x), minus vitamin A | Life Technologies | Cat# 12587010 |
| β-Mercaptoethanol | Fisher Scientific | Cat# 21985023 |
| DMEM/F12 | VWR | Cat# 45000-344 |
| Puromycin | Fisher Scientific | Cat#A1113803 |
| ROCK inhibitor Y-27632 | Fisher Scientific | Cat#50-175-998 |
| SB431542 | Stemgent | Cat#04-0010-05 |
| 96-well CYTOOplates | CYTOO | Cat#20-950-00 |
| Arena EMB CYTOOchips | CYTOO | Cat#10-021-00-18 |
| Ibidi μ-Plate 24 well black | Ibidi | Cat#82406 |
| Ibidi μ-Slide 8 well | Ibidi | Cat#80826 |
| Accutase | Fisher Scientific | NC9839010 |
| *Critical Commercial Assays* |  |  |
| DNeasy Blood and Tissue Kit | Qiagen | Cat#69504 |
| HiSpeed Plasmid Midi Kit | Qiagen | Cat#12643 |
| P3 Primary Cell 4D-Nucleofector® X Kit L | Lonza | Cat# V4XP-3024 |
| *Experimental Models: Cell Lines* |  |  |
| ESI-017 | ESI BIO | RRID:CVCL_B854 |
| ESI017 GFP-β-catenin | (Massey *et al.*, 2018) | |
| *Software and Algorithms* |  |  |
| Benchling |  | https://benchling.com/ |
| MATLAB |  | https://www.mathworks.com/products/matlab.html |
| Matlab scripts for quantifying, analyzing data | This paper | https://github.com/gbri140 |

---

**Figure S1:**
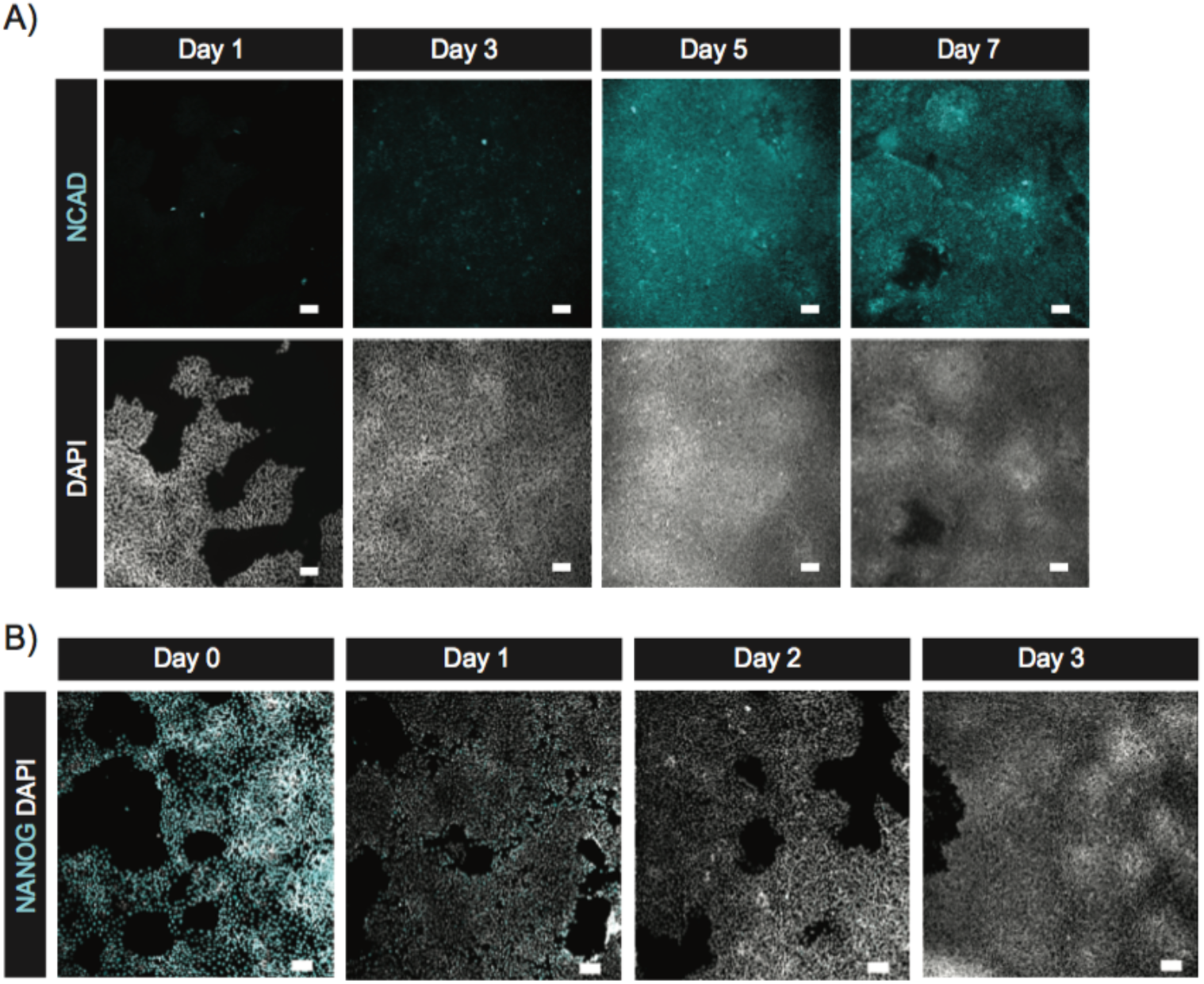
Nodal inhibition suppresses NANOG and then instructs expression of NCAD (related to Figure 1) (A-B) Representative images of cells in standard culture fixed on the indicated days during the course of Nodal inhibition and immunostained for the neural progenitor marker NCAD (A) or NANOG (B) at the conclusion of the induction. Scalebar = 100 μm.

**Figure S2:**
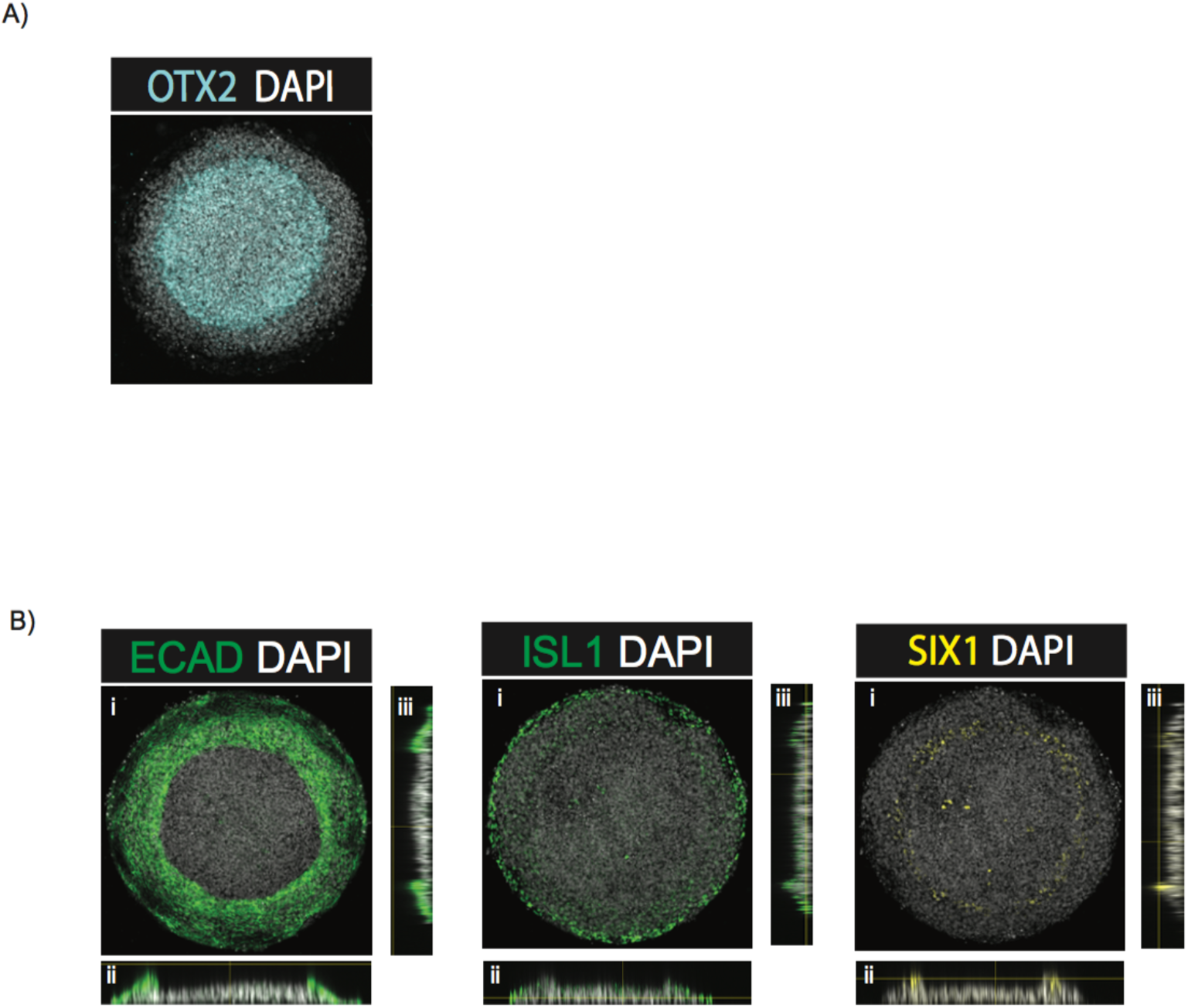
Two-step ectoderm induction protocol instructs gene expression patterns associated with anterior fates (related to Figure 2) (A-B) Representative images of following a two-step ectoderm induction protocol and immunostained for OTX2 (A), ECAD, ISL1 or SIX1 (B). Panels in B represent max projections of the indicated label in an XY (i), XZ (ii) or YZ (iii) coordinate system. Cells were initially induced for 3 days in ectoderm differentiation media and then treated with BMP4 and SB in N2B27 media for the subsequent 3 days. Colony diameter = 700 μm. Scalebar = 100 μm.

**Figure S3:**
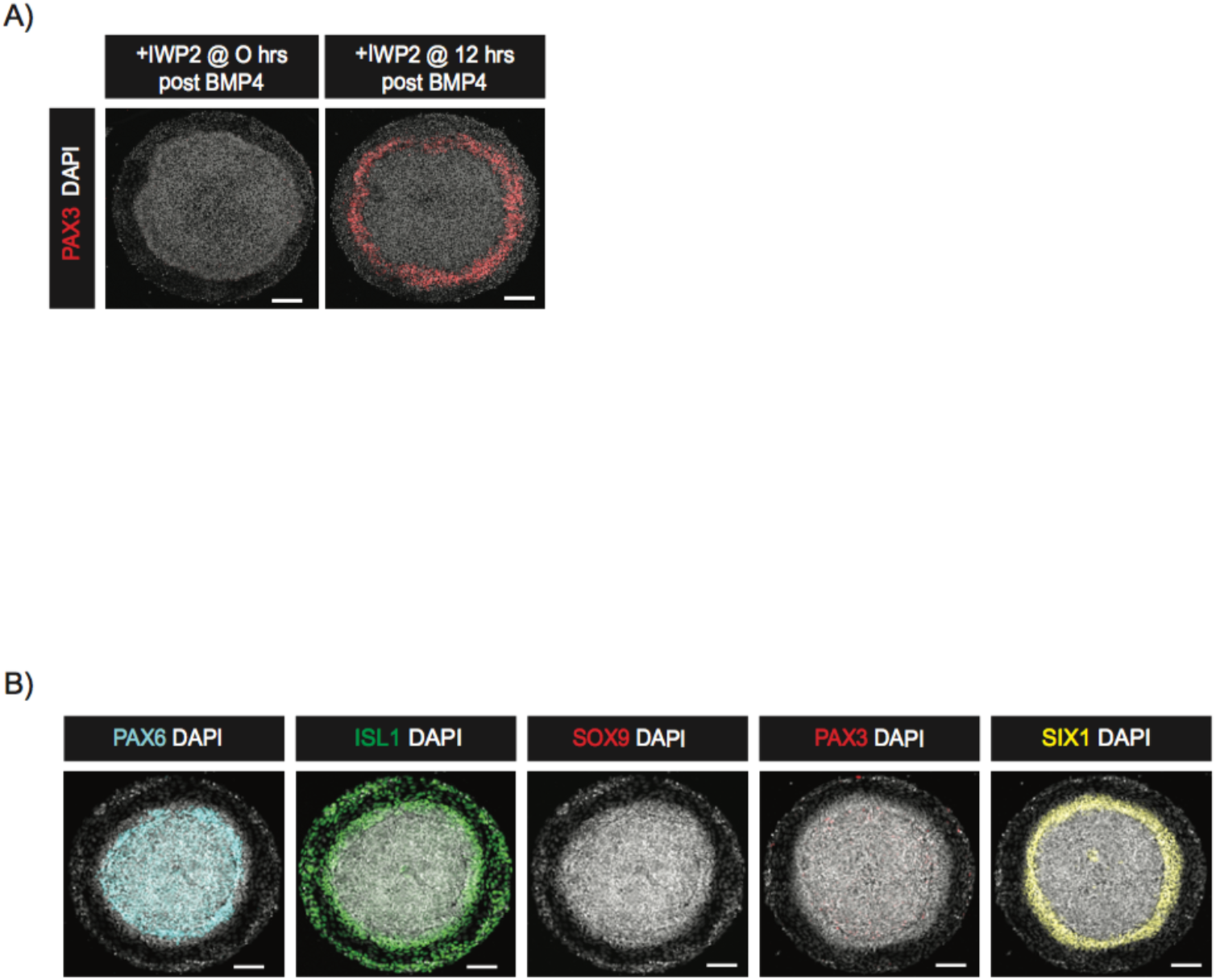
WNT inhibition before or at the time of BMP4 treatment inhibits neural crest emergence (related to figure 3) (A) Representative images of colonies immunostained for PAX3 at the conclusion of the experiment. Colonies were initially induced for 3 days in ectoderm induction media and then subsequently differentiated for 3 days in N2B27 media containing BMP4 and SB. The time between BMP4 and IWP2 addition is indicated in the banner above the corresponding image. Colony diameter = 700 μm. (B) Representative images of colonies immunostained for PAX6, ISL1, SOX9, PAX3 or SIX1 at the conclusion of induction. Colonies were initially induced for 3 days in ectoderm induction media with IWP2 and then subsequently differentiated for 3 days in N2B27 media containing BMP4, SB and IWP2. Colony diameter = 700 μm. Scalebar = 100 μm.

**Figure S4:**
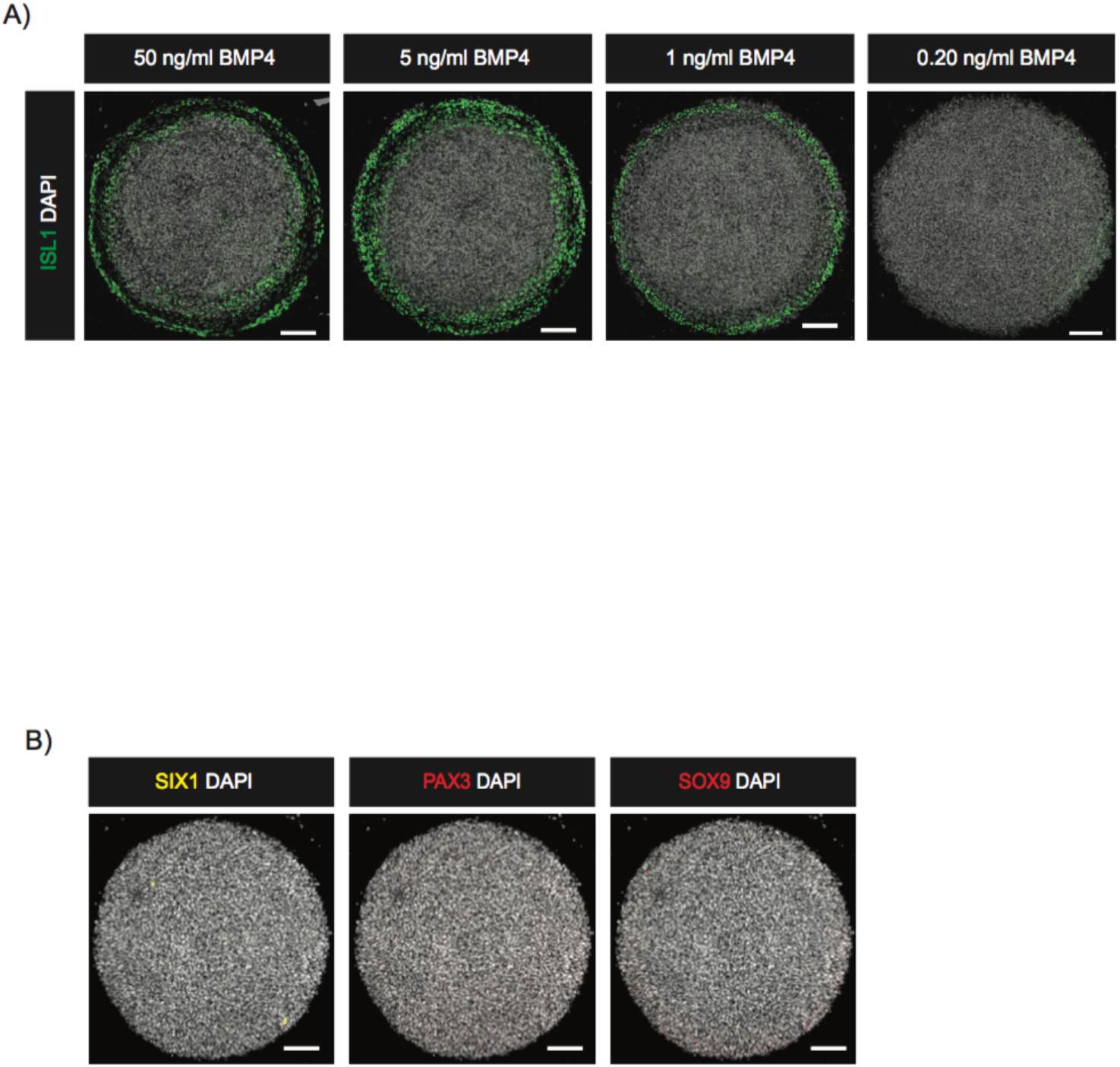
Surface ectodermal fates require a minimum concentration of BMP4 (related to Figure 4) (A) Representative images of colonies immunostained for ISL1 at the conclusion of induction. Colonies were induced with a three-step ectoderm induction protocol and patterned with the indicated concentration of BMP4. Colony diameter = 700 μm. (B) Representative images of colonies immunostained for SIX1, PAX3 or SOX9 at the conclusion of the experiment. Colonies were induced for the first 3 days in ectoderm induction media with IWP2 and then differentiated for the subsequent 3 days in N2B27 media containing 0.20 ng/ml of BMP4, IWP2 and SB. Colony diameter = 700 μm. Scalebar = 100 μm.

